# Towards whole-heart quantitative myocardial perfusion using a dual-sequence framework with multiband acceleration

**DOI:** 10.1101/2025.08.20.666501

**Authors:** Giulio Ferrazzi, Carlos Galan-Arriola, Carlos Velasco Jimeno, Carlos Real, Matteo Ghidara, Gonzalo López-Martín, Teresa Correia, Borja Ibañez, Javier Sánchez-González

**Author notes:** **Corresponding authors:** Prof. Dr. Borja Ibañez, Dr. Javier Javier Sánchez-González, Dr. Giulio Ferrazzi.

## Abstract

**Background:** 2D Quantitative Myocardial Perfusion (Qperf) MRI is limited by its inability to provide complete myocardial coverage within a heartbeat interval. This study developed and evaluated dual saturation multiband-accelerated Qperf imaging to achieve near-complete left ventricular coverage in free-breathing at an adequate in-plane spatial resolution, using a semi-automated Myocardial Blood Flow (MBF) framework for quantification deployable directly on the scanner console.

**Methods:** A dual-saturation single-band QPerf sequence was modified for multiband imaging, enabling the acquisition of 6 high-resolution myocardial slices plus Arterial Input Function (AIF) during free-breathing. The technique was evaluated in 16 sedated pigs (13 healthy and 3 with LAD occlusion) under rest conditions on a 3T MRI scanner. Additionally, two healthy pigs underwent stress imaging as well. Statistical comparisons were performed between multiband and single-band MBF values in corresponding AHA segments.

**Results:** Qualitatively, multiband QPerf provided superior left ventricular coverage and comparable image quality to single-band Qperf MBF maps, potentially enabling a more comprehensive detection of perfusion defects at rest. Quantitatively, multiband QPerf yielded lower MBF values than single-band QPerf (p < 0.01). However, Bland–Altman analysis (mean difference: −0.17 ml/min/g; 95% CI: –1.12 to 0.79 ml/min/g) and Passing–Bablok regression (intercept: –0.01 ml/min/g; 95% CI: – 0.37 to 0.28 ml/min/g) indicated that such discrepancy remained within the expected confidence intervals. Furthermore, the Passing–Bablok slope (0.88; 95% CI: 0.73–1.06) confirmed that mb-QPerf maintained sensitivity comparable to sb-QPerf in detecting perfusion changes under rest conditions. Finally, there was an overall increase in MBF values during stress vs rest conditions (average MBF Ratio 1.67 ± 0.31) when comparing healthy pigs.

**Conclusion:** Multiband-accelerated Qperf is feasible, providing improved left ventricular coverage, adequate in-plane resolution, and a semi-automated MBF quantification framework directly on the scanner console. Compared to its single-band counterpart, multiband QPerf demonstrated a more comprehensive visualization of perfusion defects and comparable sensitivity and accuracy in detecting perfusion changes at rest. Further research and clinical validation in patient populations are needed to confirm its utility in the diagnosis of coronary artery disease.

## 1. BACKGROUND

Quantitative Myocardial Perfusion (QPerf) using magnetic resonance imaging (MRI) holds potential as a radiation-free clinical protocol for the assessment of coronary artery disease (CAD) [1]. The ability to accurately quantify Myocardial Blood Flow (MBF) as opposed to the conventional, operator-dependent qualitative / semiquantitative methods [2] offers a potentially better diagnosis and prognosis tool [3].

To accurately quantify perfusion, dual-bolus and dual-sequence techniques are employed [4]. The dual-bolus method uses two gadolinium injections - one at a lower concentration - to enable precise measurement of the arterial input function (AIF) without signal saturation during the first-pass of the contrast agent. To streamline the process and avoid the complexity of administering two separate injections [5], the dual-sequence approach was developed. Unlike the dual-bolus technique, dual-sequence imaging acquires AIF data using a much shorter saturation time than conventional high-resolution perfusion sequences. This helps preventing full signal recovery of the AIF, even when using a single full contrast dose.

Traditionally, two-dimensional (2D) cartesian imaging has been the gold standard for dual-sequence rest / stress Qperf, offering high in-plane resolution for assessing ischemic damage across three to four slices [6]. While the ability to acquire data at high-resolution effectively reduces dark-rim artifacts [7,8] and helps resolving transmural perfusion defects, 2D imaging is limited by its inability to provide complete myocardial coverage during both rest and (especially) stress conditions. Whole Heart (WH) Left Ventricular (LV) coverage is nevertheless preferrable since it has been linked to a better definition of the ischemic burden [9].

Three-dimensional (3D) Cartesian k-t methods [10–14] offer the advantage of full myocardial coverage within a single cardiac phase and have also been extended to acquire multi-phase data [15]. However, this benefit comes at the expense of reduced spatial resolution, primarily due to the prolonged readout durations these methods require to freeze cardiac motion. To mitigate the issue of prolonged readout durations while maintaining sufficient in-plane resolution, non-Cartesian and hybrid 3D acquisition strategies incorporating spatial and/or temporal regularization have been proposed [16–18]. Although advances in hardware, sequence design, and reconstruction algorithms have reduced readout times considerably [19], cardiac motion remains a persistent limitation of 3D techniques, particularly during systole where the heart motion is most pronounced. Notably, systole may be the optimal cardiac phase for imaging, as the myocardium is thicker and defects are more readily detectable [15,20,21].

In parallel to 3D acquisitions, the multiband (mb) Controlled Aliasing In Parallel Imaging Results IN Higher Acceleration (CAIPIRINHA) technique has been proposed to increase anatomy coverage [22,23]. This technique simultaneously excites multiple slices and acquires them in a single readout, preserving the high in-plane resolution nature of 2D imaging, while extending coverage as it is in 3D imaging. Multiband imaging has been employed in several cardiac applications [24–26], including perfusion [20,27–33], and there is small number of pilot studies who targeted its quantification [34–36].

In this study, we present a multiband solution for Qperf imaging that: *i)* employs a dual-sequence design both during rest and stress conditions, *ii)* offers relative flexibility in selecting adequate in-plane resolution levels, *iii)* avoids temporal blurring introduced by iterative reconstruction methods with spatio-temporal regularization, *iv*) achieves near-complete LV coverage through the acquisition of 6 slices within the same heartbeat, *v)* does not require breath-holds and *vi)* provides a fully automated workflow on the scanner, enabling MBF maps to be generated directly on the console with just a few mouse clicks.

The sequence was tested on healthy and infarcted porcine models scanned under rest and stress conditions and compared against regular 3 slices Qperf [37]. Part of the work presented here has been published in proceedings of the 34th Annual Meeting of ISMRM [38].

## 2. METHODS

### 2.1 Sequence implementation

A dual-sequence single-band (sb) single-shot spoiled turbo field echo (TFE) T1 weighted Qperf cardiac perfusion sequence [37], optimized to capture within the same heartbeat both the AIF and high-resolution myocardial data, was used as the baseline sequence (Figure 1A). In this setup, the AIF module is acquired at the beginning of the cardiac cycle (dark blue module) with a short saturation time (TS) < 45ms, avoiding saturation of signal intensities within the blood pull during first-pass of the contrast agent. After AIF acquisition, three high-resolution slices (grey modules) are acquired one after another to cover the myocardium at distinct anatomical locations (i.e. basal, mid-ventricular and apical regions). Note that the sequence was designed and optimized for achieving full control of the saturation times by decoupling AIF from high-resolution acquisition modules [37].

**Figure 1.**
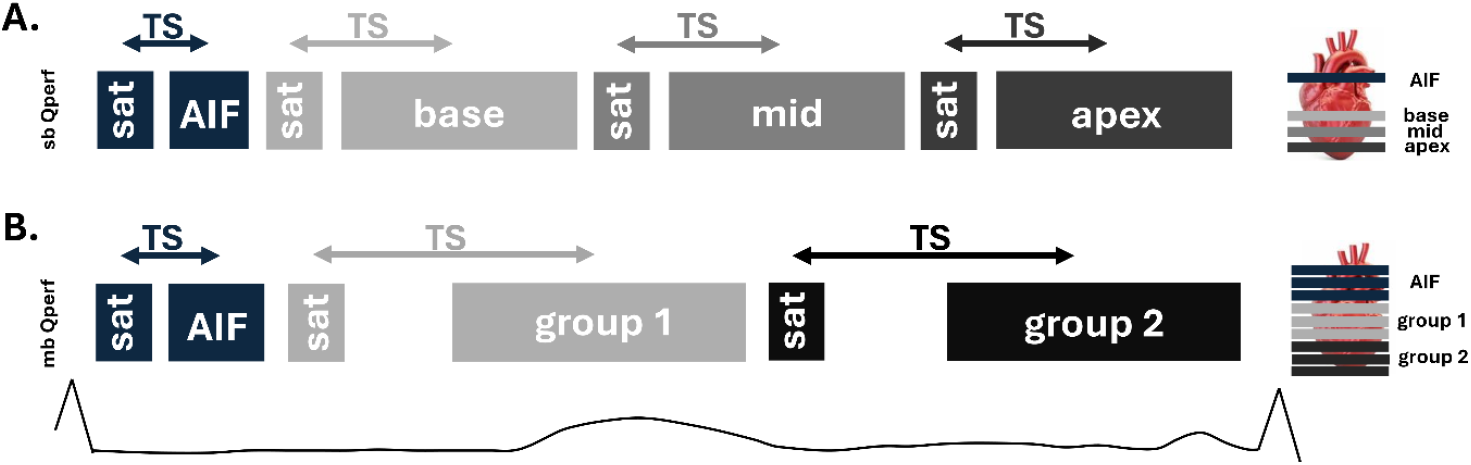
Schematic Comparison of Single-Band and Multiband Acquisition Schemes. (A) The conventional sb scheme acquires an AIF module followed by three sequential myocardial slices. (B) The proposed mb scheme acquires the AIF module followed by two multiband acquisition blocks, each simultaneously exciting three slices. The mb sequence features a longer readout duration to accommodate lower in-plane acceleration and a longer saturation time. The longer TS was intentionally chosen to reduce dark-rim artifacts [20] while maintaining compatibility with heart rates up to 110 bpm.

Building on this sb sequence, modifications were introduced to enable multiband (mb) imaging. Three triple-band (i.e. multiband 3) RF pulses were generated as a complex summation of three native sb pulses using CAIPIRINHA phase increments of 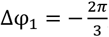, Δφ = 0 and 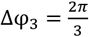 as previously described [20,25,26]. By sequentially applying these RF pulses as new k-space lines are acquired, image shifts of 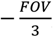, 0 and 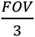 are achieved for slices 1, 2, and 3, respectively, ameliorating g-factor penalties at reconstruction [23]. Image reconstruction was performed *in-line*, where the manufacturer’s SENSE-based reconstruction workflow was adapted to process the multiband data. Note that, in this context, multiband was made compatible with conventional in-plane acceleration techniques such as SENSE and Partial Fourier.

Figure 1B shows the optimized multiband protocol, where it becomes possible to capture 6 high-resolution slices and the AIF within the same heartbeat. Please note that, for simplicity, in this prototype implementation the AIF module was multibanded as well.

### 2.2 Data acquisition

Imaging was conducted on a 3T-TX Achieva platform (Philips Healthcare, Best, The Netherlands) with a 32-channel receiver cardiac coil. Experimental procedures were performed in large porcine models (see below). All experiments were conducted in accordance with the European Directive 2010/63/EU and with Spanish national legislation (Royal Decree 53/2013). The study protocols were reviewed and approved by the competent authority (Comunidad de Madrid, CAM) under the following ethical authorizations: PROEX 24.0/25 and PROEX 120.8/22.

16 Large-White pigs (4 females) were included in the study: 13 healthy and 3 subjected to left anterior descending (LAD) permanent coronary artery occlusion. For artery occlusion, a 6 Fr guiding catheter was used to cannulate the left main coronary artery. A 0.014-inch guidewire was advanced to the distal LAD artery, and a coil was deployed in the mid-to-distal LAD using a Hunter Thrombus Aspirator (IHT, Barcelona, Spain). All animals were scanned at rest using both the mb and sb imaging protocols to establish baseline MBF values. Additionally, 2 healthy pigs underwent pharmacologically induced stress scans during separate sessions.

Pigs were sedated by intramuscular injection of ketamine (20 mg/kg), xylazine (2 mg/kg), and midazolam (0.5 mg/kg). A marginal vein in the ear was cannulated for peripheral intravenous access. Sedation was maintained by a continuous intravenous infusion of ketamine (2 mg/kg/h), xylazine (0.2 mg/kg/h) and midazolam (0.2 mg/kg/h). For stress, adenosine was administered at 0.5 mg/kg/min and, to maintain blood pressure, phenylephrine was co-infused (1-5 µg/kg/min) diluted in saline. Since swine lack significant α1-adrenergic coronary microvascular constrictor responses, phenylephrine opposes the systemic effects of adenosine, while leaving adenosine-induce coronary vasodilation unperturbed. After 1.5 minutes of the onset of adenosine administration (the peak of hyperemic effect), perfusion sequence was launched and acquired.

The sb data was collected either before or after the mb data through separate injections. Two boluses of gadoteric acid (Clariscan, GE HealthCare) were administered at least 15 minutes apart, each at a dosage of 0.5 mmol/kg and injected at a rate of 3 mL/s.

The mb and sb protocols were individually optimized to ensure high data quality and heart rates of at least 110 BPM, making them suitable for stress imaging (Figure 1). The mb protocol had the following parameters; short/long TS = 35/135ms, in-plane resolution = 2.6×2.6mm^2^, slice thickness = 9mm, slice gap = 1mm, TE/TR = 1.06/2.3ms, flip angle α = 15°, shot duration (high-resolution data) = 159.9ms, mb factor = 3, SENSE factor = 1.35, Partial Fourier = 0.81 and number of reconstructed slices = 9. The sb protocol was acquired with a short/long TS = 30/85ms, in-plane resolution = 2.6×2.6mm^2^, slice thickness = 10mm, slice gap = 0mm, TE/TR = 1.04/2.26ms, flip angle α=15°, shot duration (high-resolution data) = 105ms, SENSE factor = 2, Partial Fourier = 0.75 and number of reconstructed slices = 4.

The following elements were common to all mb and sb scans and imaging protocols:

i. a reverse linear k-space ordering was used, as it offers improved Point Spread Function characteristics when combined with Partial Fourier acquisition [37];
ii. integrated proton density (PD) images, acquired without a saturation pulse during the first heartbeat for normalization purposes [37];
iii. two baseline MOLLI T1 maps acquired before each contrast injection to enable accurate conversion of contrast agent concentration [39]; and
iv. hematocrit values recorded prior to the scanning sessions for conversions purposes [37].

### 2.3 Motion correction, AIF selection and MBF quantification

MBF quantification was performed entirely on the scanner. Following image acquisition, motion correction was applied using the Fast Elastic Image Registration (FEIR) algorithm [40], applied across all dynamic frames for both sb and mb datasets. The resulting motion-corrected images were subsequently used for perfusion analysis.

For sb acquisitions, the AIF was determined from a designated low-resolution aortic slice. A region of interest (ROI) was manually drawn within the aortic outflow tract, and the mean signal intensity was recorded at each time point to generate the AIF curve. For mb data, the AIF was extracted from the first of the three simultaneously acquired AIF slices.

MBF quantification was performed following the framework described in [37]. The normalized signal intensity curves, 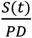, were first converted into contrast agent concentration-time curves using pre-contrast myocardial and blood T1 values obtained from MOLLI. Blood flow was then estimated through a deconvolution process based on the Tofts model [41]. Finally, it was converted into MBF values using the recorded hematocrit.

A detailed description of this MBF quantification methodology is provided in [37].

### 2.4 Myocardial Segmentation and AHA Analysis Tool

A custom graphical user interface (GUI) was developed in Python to facilitate the segmentation and analysis of cardiac MBF maps.

Myocardial endocardial and epicardial borders were manually delineated on a slice-by-slice basis. For each slice, a single anatomical reference point, corresponding to the anterior right ventricular (RV) insertion, was identified in accordance to [42]. This reference point was used to enable automatic segmentation of the myocardial tissue into the American Heart Association (AHA) 16-segment model. As in the mb case there are more than 3 slices, in addition, each slice was manually annotated as belonging to ether the apical, mid, or basal region of the heart. Pixel-wise statistics - including mean and standard deviation were computed within each of the 4 or 6 segments defined per slice. These per-slice statistics were subsequently averaged across regions to generate the full set of 16 AHA segments across the entire image stack.

### 2.5 Statistical Analysis

The data from the rest scans underwent statistical testing.

Statistical comparisons between sb and mb AHA segments across all animals were performed. The analysis included the following steps: *(i)* assessment of data normality for the sb and mb groups separately using the Shapiro–Wilk test; *(ii)* paired t-tests across distributions (if both distributions were normal) or the Wilcoxon signed-rank tests (if normality was rejected for one or both distributions). In all tests, a p-value lower than 0.01 was considered statistically significant. Additionally, agreement between sb and mb pairs was evaluated using Bland–Altman plots using the 95% confidence intervals (CI). Passing–Bablok regression, with estimation of slope and intercept along with their corresponding 95% CI was performed as well.

## 3. RESULTS

Figures 2 and 3 present representative cases comparing sb and mb-Qperf maps in animals with LAD occlusion. In the case shown in Figure 2, regions with lower perfusion are visible within the expected ischemic segments for both sequences. While MBF values are comparable between sb and mb within these regions, the MBF values recorded in the surrounding healthy myocardium are higher in the mb maps. In the case shown in Figure 3, both methods record similar MBF in healthy segments. The visualization of the perfusion defect differs, with the mb map depicting an ischemic territory that spans the entire myocardium from base to apex.

**Figure 2.**
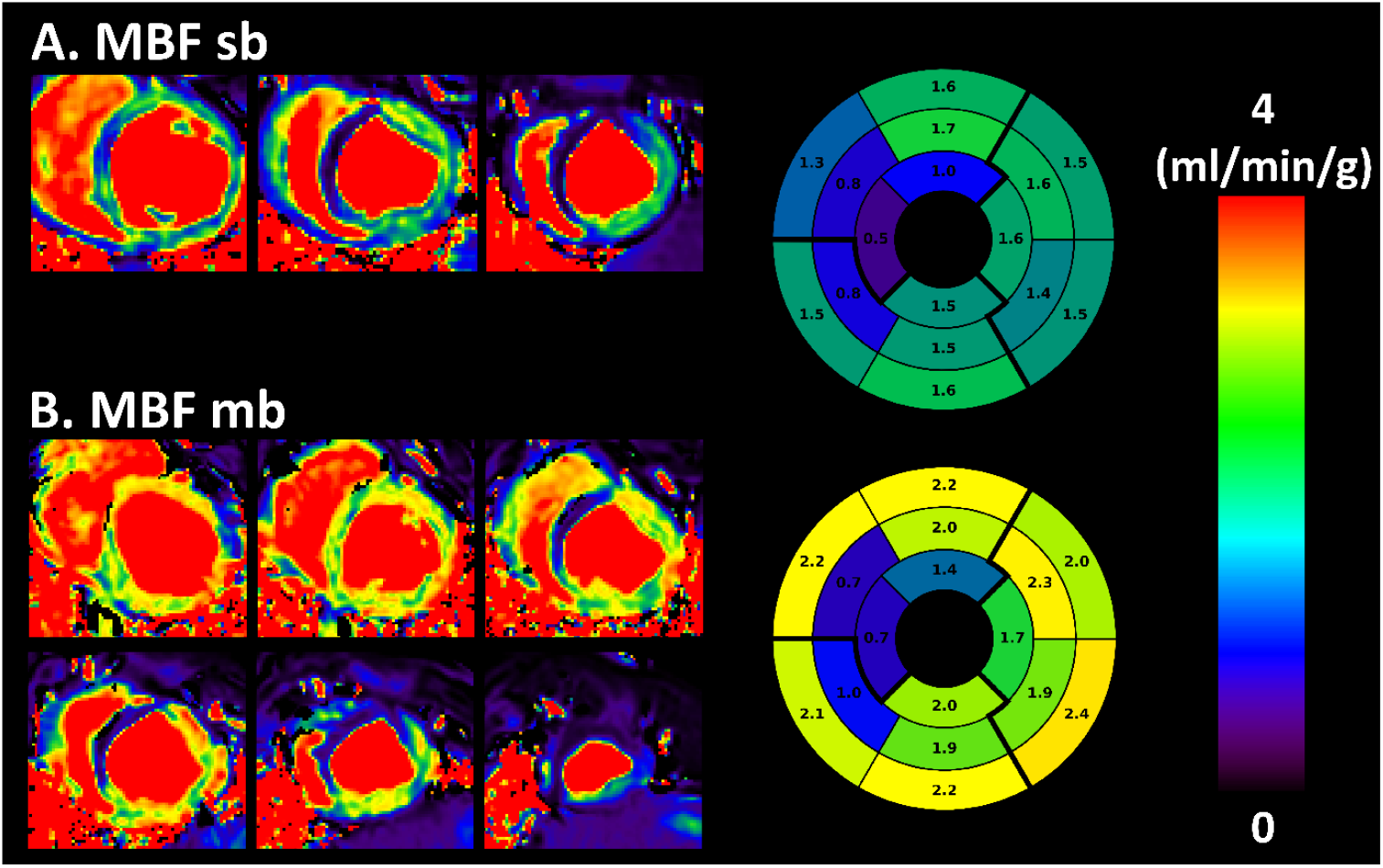
Representative MBF Maps in the first animal. Comparison of MBF maps and corresponding bull’s-eye plots from single-band (panel A in each figure) and multiband (panel B in each figure) acquisitions. Hypo-perfused regions are clearly visualized with both techniques, demonstrating comparable qualitative performance for detecting perfusion defects.

**Figures 3.**
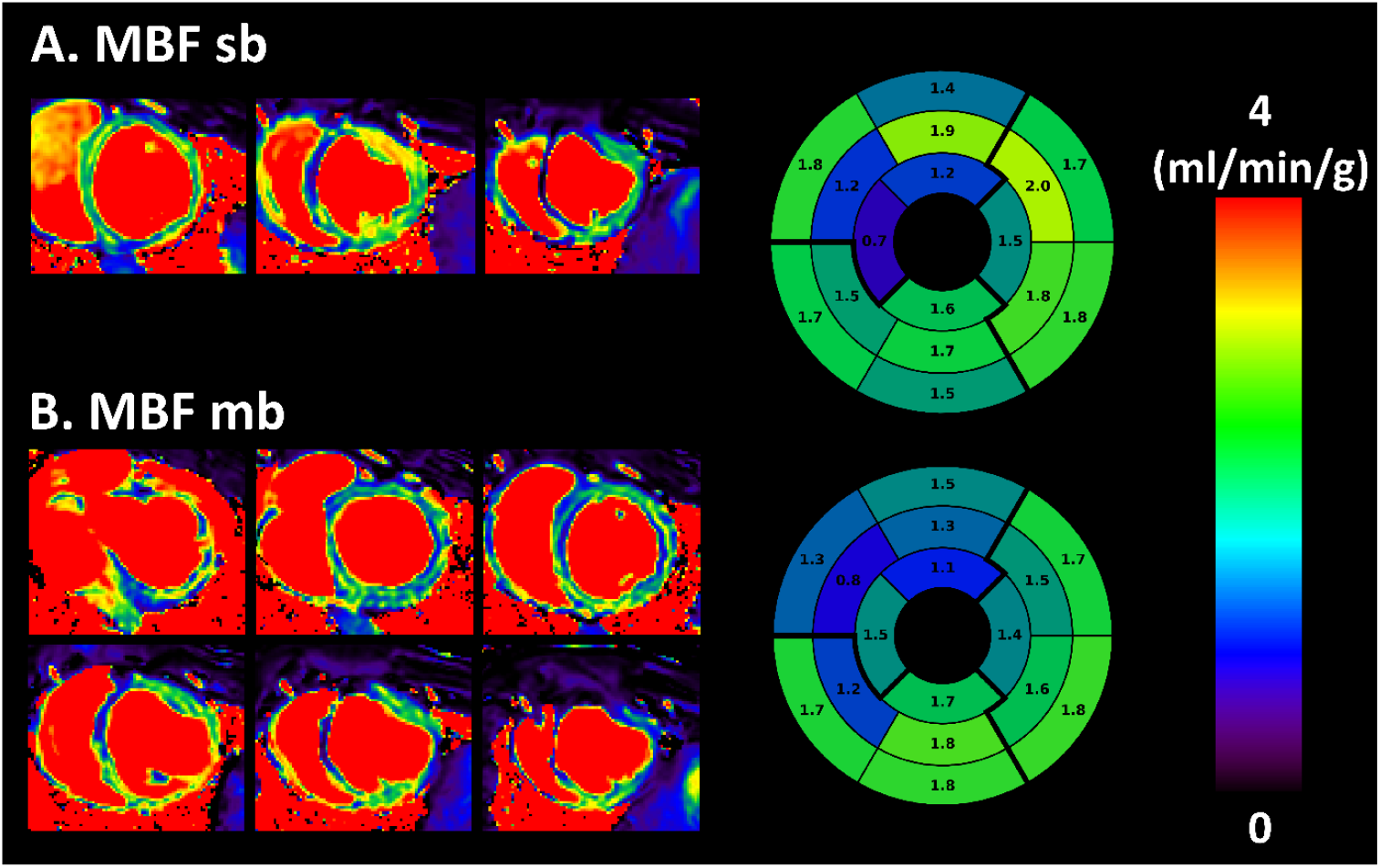
Representative MBF Maps in the second animal. Same as Figure 2 in the second animal.

For the statistical tests, both MBF sb and mb values demonstrated a statistically significant deviation from normality (*p* < 0.01). The Wilcoxon signed-rank test revealed a statistically significant difference between the sb and mb distributions (*p* < 0.01). Figure 4A shows Bland-Altman plots comparing MBF values between corresponding segments in the sb and mb datasets. No apparent trend is observed, with mb generally displaying lower values than sb (mean difference: –0.17 ml/min/g). The 95% CI range (–1.12 ml/min/g to 0.79 ml/min/g) includes the zero line, indicating the absence of a systematic bias within the limits of agreement. Figure 4B presents the Passing-Bablok regression analysis, which revealed a slope of 0.88 and a 95% CI containing 1 (CI: 0.73–1.06), suggesting that sb and mb have comparable sensitivity to perfusion changes within the limits of the agreement. Moreover, the 95% CI for the intercept (–0.01 ml/min/g; CI: –0.37 ml/min/g to 0.28 ml/min/g) did include zero, indicating again the absence of a systematic bias between the two measurements.

**Figure 4.**
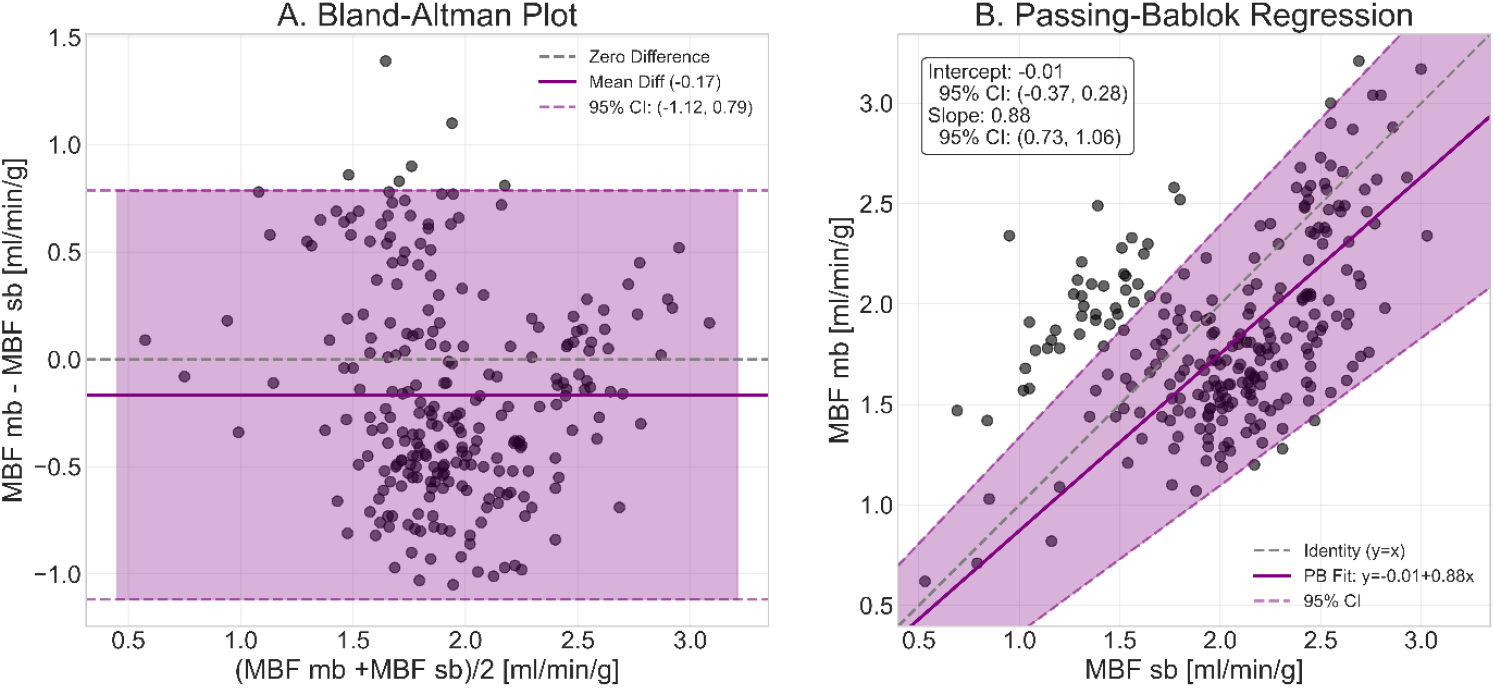
Quantitative Comparison and Agreement of Single-Band and Multiband MBF. (A) Bland–Altman plot comparing segmental MBF values from the sb and mb acquisitions. (B) Passing–Bablok regression analysis.

Figure 5 (left) displays MBF maps from stress and rest scans in two healthy pigs, acquired in separate sessions. On the right, the MBF Ratio (MBFR) - calculated as the segment-by-segment ratio of the individual AHA plots - is shown. There is an overall increase in MBF during stress vs rest conditions, with an average MBFR of 1.67 ± 0.31 across all segments and animals.

**Figure 5.**
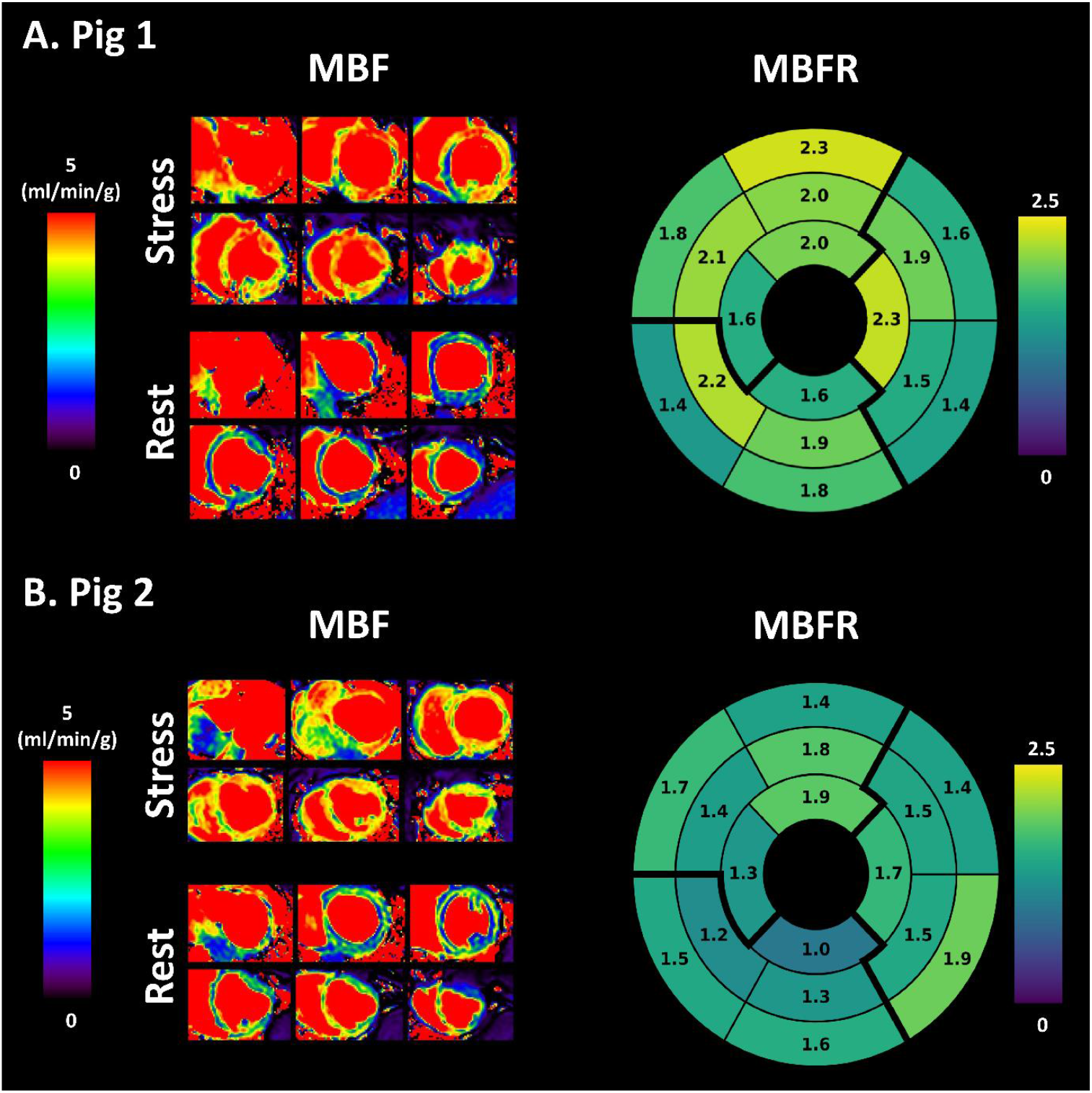
Myocardial Perfusion Reserve (MPFR) Assessment. Comparison of rest and stress MBF maps and the resulting calculated MPFR maps for (A) the first subject and (B) the second subject. The multiband sequence effectively captures the increase in perfusion during stress relative to rest.

## 4. DISCUSSION

This study introduces and validates a novel multiband quantitative myocardial perfusion technique in a porcine model. The primary objective was to address the restricted cardiac coverage of 2D single-band approaches and the motion sensitivity and spatial resolution challenges inherent in 3D techniques. Our proposed method provides near-complete LV coverage with adequate in-plane resolution during free-breathing and incorporates a semi-automated workflow for on-scanner MBF quantification. To evaluate its performance, the technique was validated against the sb-QPerf reference standard. Its capabilities were further demonstrated through representative stress imaging in healthy pigs and by characterizing large, established infarcts at rest in three separate cases.

The application of multiband acceleration to quantitative myocardial perfusion imaging is an emerging field of research and the first studies are starting to appear [34–36]. From a clinical standpoint, Nazir et al. [34] demonstrated high diagnostic accuracy for determining significant epicardial CAD with a six-slice quantitative cartesian multiband protocol, validating it against fractional flow reserve and quantitative coronary angiography. While this represents a landmark clinical evaluation, the study lacked a direct quantitative validation of the multiband MBF values against a single-band reference. Furthermore, the omission of critical technical details, such as AIF implementation / placement may hinder reproducibility. The 2D multiband radial sequence pioneered by Huang et al. [35] represents a significant technical advancement in the field. Their framework offers key advantages, including AIF sampling from the initial radial spokes of the readout and the potential for high-resolution, cardiac-phase-resolved reconstructions at systole and diastole. Despite these promising features, the full potential for expanded coverage was not assessed, as the validation was limited to a three-slice protocol. Finally, Tian et al. [36] employed an interleaved radial multiband approach to achieve whole-heart, ungated, free-breathing quantitative MBF maps without magnetization preparation. However, this rather complex framework has limitations as it requires the administration of two separate contrast injections for quantification and may not guarantee steady-state conditions in the outermost slices due to cardiac motion [36]. As a crucial first step, in this study we opted for a Cartesian sampling multiband scheme. This choice was motivated by our goal to deliver a solution that could be seamlessly integrated with existing on-scanner reconstruction software. By prioritizing on-scanner compatibility, we have established a robust, clinically-translatable framework. Having validated the quantitative accuracy of the technique, our next step is to extend this framework to radial [30] / spiral [29] under-sampling to improve efficiency.

In this study, which focused on large, non-reperfused infarcts, significant differences in diagnostic outcome between sb-QPerf and mb-QPerf were not observed (Figures 2 and 3). However, the enhanced LV coverage of the proposed mb-QPerf technique (6 slices vs. 3 slices with sb) represents a key technical advantage. We hypothesize that this comprehensive coverage will become critically important for the accurate assessment of smaller or more subtle perfusion defects, particularly in challenging regions such as the apex, which may be missed or only partially covered by sb acquisitions. Furthermore, acquiring multiple, closely spaced slices with mb offers data redundancy that may improve segmentation accuracy in difficult cases, such as in thinned myocardial walls. Validating the clinical impact of mb-QPerf in these specific scenarios remains an important objective for future investigation.

This study has also limitations. While meeting published guidelines [43] achieving an in-plane resolution < 3mm isotropic, the readout time of the multiband protocol was slightly longer (159.9ms) than the 100-125ms range recommended for freezing cardiac motion [43]. These parameters are a consequence of a significant overall acceleration factor of 5 (MB=3, SENSE=1.35, Partial Fourier=0.81), which pushes the boundaries of current parallel imaging capabilities. During optimization, further shortening the readout via increased partial Fourier and higher SENSE factors combined with multiband led to excessive g-factor noise amplification. Thus, the current configuration represents a carefully balanced compromise in which a pragmatic approach was adopted, prioritizing fast, on-scanner reconstruction without temporal regularization, as its impact on quantification is not well understood and depends on several factors [13]. Notably, these results were achieved at 3T which may prove too aggressive for lower field strengths such as 1.5T. Another limitation of this study is that stress imaging was performed in only two healthy animals. Although validation under rest conditions offers a valuable initial proof of concept, clinical decision-making predominantly relies on stress imaging. As such, the true clinical utility of the proposed approach must be evaluated under stress conditions. Future studies will aim to address this by incorporating LAD occlusion during stress imaging in animal models. Ultimately, translation to human studies will also be necessary to fully validate the method’s applicability in clinical practice. Notably, the ability of mb-QPerf to detect large perfusion defects at rest similarly to sb-QPerf already offers compelling evidence that whole-heart imaging is clinically valuable. These early results strongly suggest that mb-QPerf has the potential to significantly enhance ischemia detection and quantification - a hypothesis that ongoing and future studies under stress conditions are now well-positioned to confirm.

## 5. CONCLUSION

This study validates multiband CAIPIRINHA-accelerated dual-sequence QPerf as a robust technique for comprehensive myocardial perfusion imaging. The method successfully achieves complete left ventricular coverage and adequate resolution within a free-breathing acquisition, all processed by an automated on-scanner quantification framework. Visually, mb-QPerf delivers image quality comparable to the standard single-band technique, with the enhanced coverage offering the potential for improved defect detection. While mb-QPerf showed a slight statistical significant underestimation of MBF values, this bias was well within the limits set by the confidence internal and did not compromise the sensitivity to perfusion changes at rest.

